# Sharing the trail: recreation effects on bear behaviour in a Canadian Rocky Mountain Park

**DOI:** 10.64898/2026.04.02.714576

**Authors:** Alexandra Dimitriou, Kaitlyn M. Gaynor, Sarah Benson-Amram, Melanie Percy, A. Cole Burton

## Abstract

Humans are profoundly reshaping the natural world. These changes are giving rise to complex and mutually risky dynamics between people and large carnivores. In protected areas across North America, bears (*Ursus sp*.) face rapidly rising recreation pressures that can alter their use of the landscape, either displacing them from high-quality habitats or drawing them into human-wildlife conflicts through habituation or attraction to anthropogenic resources. However, disentangling responses to recreation from other drivers can be difficult because human activity covaries with environmental and seasonal processes that also shape bear activity. We leveraged the partial closure of the popular Berg Lake Trail in Mount Robson Park, British Columbia, Canada, to investigate whether black (*Ursus americanus*) and grizzly bears (*Ursus arctos*) showed fear, attraction or neutral behavioural responses to varying recreation levels across multiple spatiotemporal scales. To understand both anticipatory responses to predictable patterns of human activity, and reactive responses to hiker events, we used detections from 43 camera traps over two years (July 2023-June 2025). We compared weekly habitat use, daily activity patterns, and direct responses to hikers (using Avoidance-Attraction Ratios; AARs) among camera sites and between open and closed sections of the trail. Our results revealed that both bear species exhibited patterns consistent with fear responses, while some black bear behaviours were also consistent with attraction responses. Both kinds of responses reflect anticipatory strategies rather than reactionary behaviours (i.e., no AAR effect). Neither species avoided recreation spatially at the weekly scale: black bears were detected more at site-weeks with greater recreation intensity, while grizzly bears were consistently detected more at sites closer to hiking trails. However, both species used daily temporal partitioning to avoid direct encounters with humans. These findings demonstrate scope for human-bear coexistence when recreation levels are managed to be moderate and predictable, and bears have sufficient space to segregate from humans during peak times. Thus, successful coexistence will hinge on co-adaptation by both bears and people. Understanding how recreation influences bear behaviour, and the spatiotemporal scale at which that occurs, is critical for guiding effective adaptive management aimed at fostering human-bear coexistence in high-traffic protected areas.

---

Coexisting with large carnivores has long been one of humanity’s greatest conservation challenges. As human activity expands into every corner of the natural world, accelerating the loss of Earth’s remaining wild spaces, interactions between people and large carnivores are shifting in ways that increasingly challenge the delicate balance of coexistence (Bautista et al., 2017; Bombieri et al., 2019, 2023; Lamb et al., 2020). Across North America, Indigenous peoples have stewarded lands and wildlife through relationships rooted in kincentricity and reciprocity since time immemorial (Bhattacharyya & Slocombe, 2017; Martinez et al., 2023), but European colonization has led to expanding urban development, resource extraction, and other human activities that are now encroaching on the last tracts of large carnivore habitat, including in protected areas (PAs). These pressures fragment the large landscapes that carnivores need to persist, reducing functional connectivity and thereby, creating vulnerable, isolated sub-populations and driving their ongoing declines. This fragmentation also pushes carnivores into human-dominated spaces more often, increasing the frequency and intensity of human-wildlife conflicts (Bombieri et al., 2023; Gaynor & Green, 2026; Miller et al., 2020; Palm et al., 2025; Penteriani et al., 2016; Wolf & Ripple, 2017). In the context of today’s biodiversity crisis, large carnivores remain among the most imperilled species, despite their critically important ecological roles and cultural values (Burton et al., 2024; Ceballos & Ehrlich, 2002; Clark et al., 2021; Morrison et al., 2007; Ripple et al., 2014).

While large carnivores evolved at the top of food webs, humans now act as a dominant predator in many shared landscapes, profoundly influencing the behaviour and movement of other species (Darimont et al., 2015; Dorresteijn et al., 2015; Estes et al., 2011; Gaynor et al., 2019; Smith et al., 2017). Many species perceive humans as risky and may alter their behaviour to reduce interactions with them both temporally, by shifting activity to times when humans are less active, and spatially, by avoiding areas with high human presence altogether (e.g., hiking trails or campgrounds; Frid & Dill, 2002; Procko et al., 2023; Wang et al., 2015). This response is often most pronounced in large carnivores, which are less tolerated by people and for whom encounters with humans can be—and across evolutionary timescales, have been—dangerous and often lethal for both parties (Burton et al., 2024; Lamb et al., 2020; Ripple et al., 2014; Zenth et al., 2025). The risk disturbance hypothesis describes the way that animals perceive human presence as a sort of predation risk and respond accordingly (Frid & Dill, 2002; Peters & Otis, 2005). Even when engaging in non-consumptive and quiet activities such as hiking, which are generally considered to be ‘low impact,’ humans can still disrupt natural behaviours and elicit antipredator responses (e.g., fleeing, increased vigilance; Frid & Dill, 2002; Gaynor et al., 2025; Gaynor & Green, 2024). While innate fear of a perceived predator typically causes reactive responses to visual, auditory or olfactory cues that an animal associates with risk (e.g., fight or flight responses such as black bears climbing trees to avoid threats; Herrero, 1972), animals can learn to predict the temporal and spatial patterns of human activity to minimize stressful encounters (Frid & Dill, 2002; Gaynor & Green, 2024; Wheat & Wilmers, 2016; Zeller et al., 2024). Anticipatory behaviours demonstrate associative learning and behavioural plasticity as animals integrate their experiences into their decision-making (Gaynor & Green, 2024; Petrullo et al., 2024; Smith et al., 2021; Zenth et al., 2025).

When animals have had sufficient experience with humans, they can employ proactive, anticipatory strategies to reduce the frequency and intensity of encounters with humans (Gaynor et al., 2025). Avoidance allows them to maintain a lower baseline stress level without sacrificing key activities, such as foraging, thereby maintaining fitness (Wheat & Wilmers, 2016; Crupi, 2004). In particular, when human activity is both frequent and predictable, animals can adjust their use of the landscape in time and space, based on those patterns of human activity (Lima & Dill, 1990; Palmer et al., 2022, 2023). Anticipatory avoidance, under a dynamic landscape of fear framework, is conducive to coexistence by allowing wildlife to reduce risky encounters with humans. However, when the costs of avoidance outweigh the benefits, or after many benign interactions with humans, animals can become habituated. Habituation is a decrease in the strength of that fear response after repeated presentations of a stimulus when there is no benefit in responding to the stimulus (Rankin et al., 2009). Repeated benign interactions with humans allow animals to reasonably predict that humans are unlikely to harm them, and thus operate at lower levels of vigilance (Engelhardt & Weladji, 2011; Gunther & Bramblett, 2018; Samia et al., 2015). Carnivores may become habituated to recreationists following repeated, non-negative encounters, which can both facilitate and result from access to anthropogenic resource subsidies such as food (e.g., trash left behind by hikers; Beckmann & Berger, 2003; Herrero & Higgins, 2003; Newsome et al., 2015; Newsome & Van Eeden, 2017) or protection from intraguild competitors and dominant conspecifics (e.g., to avoid sexually selected infanticide; Palombit, 2015; Steyaert et al., 2016). Habituation becomes problematic when human-wildlife conflicts arise due to decreased wariness in large carnivores, coupled with overcrowded recreational spaces and irresponsible behaviours from recreationists, such as wildlife feeding or poor attractant storage (Bombieri et al., 2021; Geffroy et al., 2015; Herrero et al., 2005; Morales-González et al., 2020; Orams, 2002).

Bears (*Ursus sp.*) are highly adaptive carnivores whose behavioural plasticity allows them to persist in dynamic, human-modified landscapes. Although they are large-bodied with substantial space and energy requirements, traits typically associated with elevated conflict risk, their behavioural flexibility enables them to exploit human-dominated areas (Bombieri et al., 2023; Burton et al., 2024; Lamb et al., 2020; Vargas Soto et al., 2022). This adaptability can become maladaptive when it increases bears’ exposure to anthropogenic foods and thereby elevates the likelihood of human-bear conflict (Gunther et al., 2004; Herrero et al., 2005; Lamb et al., 2020). While risk-tolerance and boldness are genetically influenced, social learning and lived experience also shape bear behaviour, with intergenerational transfer reinforcing patterns of avoidance or habituation at the population level over time (Blair et al., 2020; Hopkins, 2013; Morehouse et al., 2016; Tuomainen & Candolin, 2011). Consequently, bears are one of the most conflict-prone taxa globally (Bombieri et al., 2023; Lamb et al., 2020; Penteriani et al., 2020).

Human-bear conflict is a key management concern in the Canadian Rocky Mountain Parks, where high levels of nature-based tourism and abundant bear populations coincide (Chamberlain et al., 2012; Ladle et al., 2018; Orr, 2014). This overlap underscores the need to better understand the mechanisms driving behavioural adaptation to humans in order to generate preventative management strategies that protect wildlife and recreationists. To address this need, we used camera trap sampling across a gradient of human activity, in both open and closed sections of a popular trail, to test how black bears (*Ursus americanus*) and grizzly bears (*Ursus arctos*) responded to the presence of recreationists. We tested three alternative hypotheses for bear responses to recreation activity. First, the fear hypothesis posits that bears perceive humans as a source of mortality risk, and thus their behaviour reflects fear and avoidance of people (Burton et al., 2024; Gaynor et al., 2019; Palmer et al., 2022). Under this hypothesis, we predict that bears will either avoid areas where they expect high levels of human activity or adjust the timing of their activity away from the timing of human activity when they do use these areas. Alternatively, under the attraction hypothesis, bears may not show strong avoidance of people but rather may be attracted by potential benefits such as access to food resources or refuge from dominant conspecifics. As such, bears would be attracted to areas and times where they expect high levels of human activity (Albert & Bowyer, 1991; Greenleaf et al., 2009). In the case of black bears (*Ursus americanus*) and female grizzlies with cubs, this behaviour may also reflect a human shield effect, where proximity to people offers protection from dominant conspecifics (Edwards, 2023; Machutchon et al., 1998; Nevin & Gilbert, 2005). Within the attraction hypothesis are two sub-hypotheses. First, under the food-motivated sub-hypothesis, if bears are seeking anthropogenic subsidies, we would expect bears to be detected more during high recreation times and areas. Second, the human shield sub-hypothesis predicts that black bears will use hikers as protection from grizzly bears, and thus that black bear activity will be higher, and grizzly bear activity lower, in areas or times of high recreation. Although the fear and attraction hypotheses describe opposing responses, they are not mutually exclusive and may operate simultaneously across spatiotemporal scales. Lastly, bears may not strongly fear nor be attracted to people and instead perceive them as a benign presence (e.g., via habituation). Under the neutrality hypothesis, use of the landscape in time and space would not be influenced by human activity, or the associated risk and rewards may cancel one another out, leading to no observable response to recreationists at any scale.

## STUDY AREA

The Berg Lake Trail is located in Mount Robson Park (MRP), British Columbia, Canada, which spans an area of 2,253 km^2^ within the Canadian Rocky Mountain Parks UNESCO World Heritage Site and has a rich history of mountaineering and hiking dating back to its establishment in 1913 (BC Parks, n.d.). Therefore, recreationists and bears have been sharing this landscape for over 100 years. MRP is home to two species of bear: black bear and grizzly bear, as well as several prey species (e.g., white-tailed [*Odocoileus virginianus*] and mule deer [*Odocoileus hemionus*], elk [*Cervus canadensis*], mountain goat [*Oreamnos americanus*], hoary marmot [*Marmota caligata*] and small rodents) and competitors (e.g., cougars [*Puma concolor*], coyotes [*Canis latrans*], wolves [*Canis lupus*]; Figure S1 in Supporting Information). While black bears are numerous and not of conservation concern, grizzly bears are listed as special concern both provincially and federally (Province of British Columbia, n.d.; SARA, 2002). MRP’s most popular trail, the Berg Lake Trail, is 23 km in length and is connected to several adjacent routes (e.g., Hargreaves Lake, Toboggan Falls, Mumm Basin, and Snowbird Pass Routes), traversing a diverse mosaic of ecosystems from inland cedar-hemlock rainforests to high alpine tundra. Low-impact, non-motorized activities like hiking and backcountry camping are the main forms of human activity along the Berg Lake Trail.

In a typical year, the Berg Lake Trail attracts over 115,000 recreationists, with up to 1,000 people using the trail each day during peak season (Environment and Climate Change Strategy, 2023). However, in 2021, an extreme heatwave triggered accelerated glacial melt, causing the Robson River to reroute and flood the trail, prompting immediate evacuation and closure (Baum et al., 2026; BC Parks, 2022). The provincial park agency (BC Parks) then initiated a phased reopening beginning in 2023 and ending with the full reopening of the trail in June 2025 (Figure S2). While the open section of the trail was fully open to the public without restrictions, the closed section of the trail had significantly lower human presence, limited to the research team, park staff, trail construction crews, and occasional trespassing hikers (Figure 2). This closed section functions as a quasi-experimental control, allowing us to compare two distinct forms of human-wildlife interactions: frequent interactions in the open portion, where there is a regular flow of humans, and bears may expect a high risk of human interaction during the summer months, and infrequent interactions in the closed portion, where the risk of interacting with humans is far lower and more unpredictable.

## METHODS

### Study design

In July 2023, we deployed 42 Reconyx HyperFire2 (Reconyx, Holmen, USA) camera traps (CTs) on and around the Berg Lake Trail and connected trails (Figure 1). CTs were fastened to trees or PVC pipe structures approximately 0.5–1 m above the ground, oriented perpendicular to a target area of expected animal or human travel (e.g., hiking or game trails) at a distance of 3–5 m. CTs were programmed to capture motion-triggered images, as well as a single daily time-lapse image at noon to monitor functionality. We randomly selected CT deployment locations within two strata: on-trail or off-trail. The on-trail stratum consisted of CTs placed directly facing hiking trails, with a minimum spacing of 500 m between sites. The off-trail stratum consisted of random locations situated 250–500 m from the nearest trail. At each off-trail location, the camera was positioned at a nearby game trail or animal sign within a 100-meter radius of the randomly generated point. When this was not feasible due to terrain constraints (e.g., water bodies, glaciers, cliffs), the CT was placed at the nearest suitable site. Initially, we deployed 24 CTs on-trail and 18 off-trail; however, as the trail was repaired and rebuilt to avoid future flooding events, some off-trail CTs became on-trail CTs as new trail sections were built in front of them. These cases are labelled as “Mixed” trail status or closure status in Figure 1, to signify that the state of the trail section (open/closed) or CT site (in-trail/off-trail) was temporally variable.

**Figure 1.**
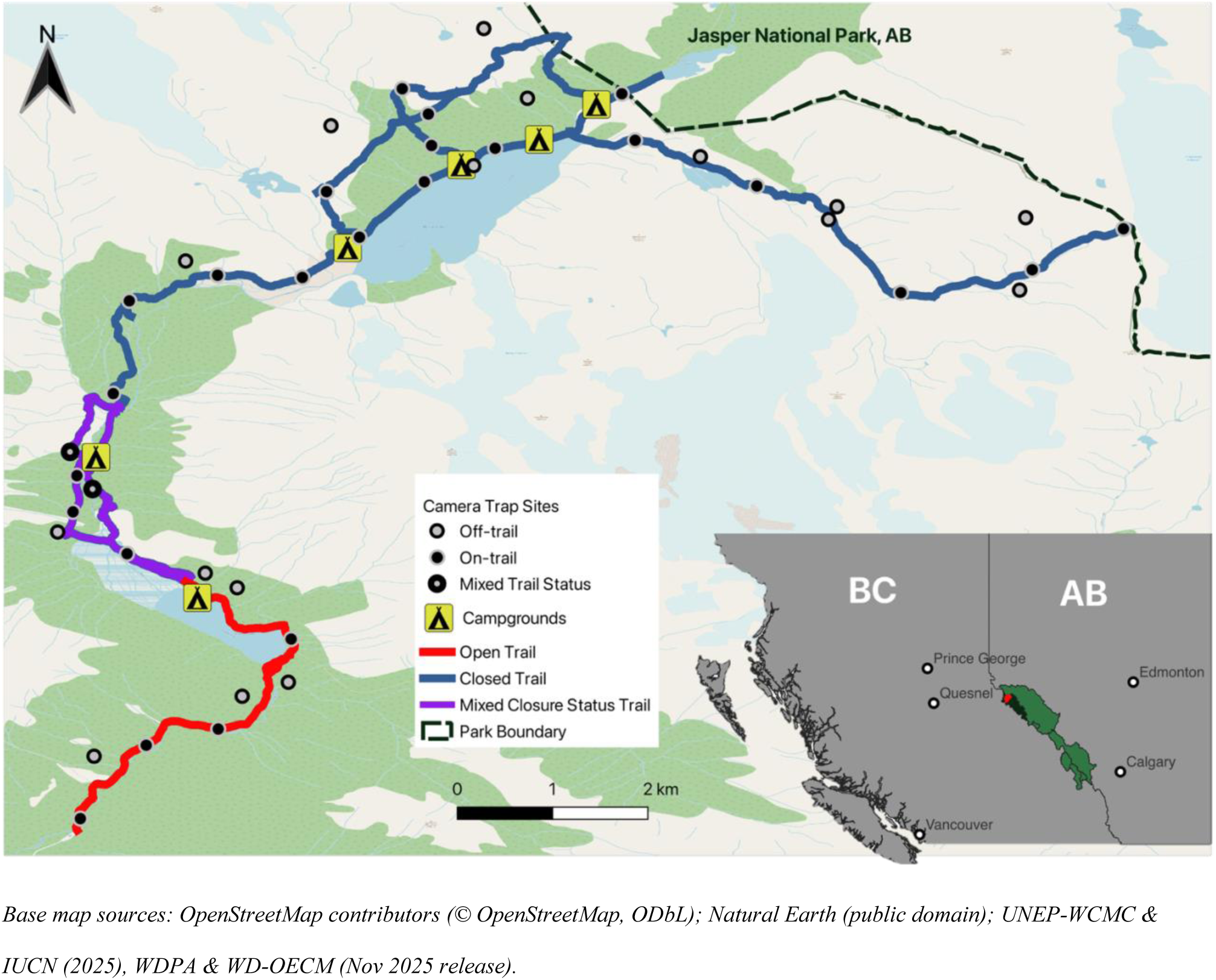
Map of camera trap stations in Mount Robson Park. Red trails were open to the public throughout the study period (July 2023-July 2025), blue trails were closed. Mixed closure trail sections (purple) were either open for a period or built during the study period. Grey points represent off-trail CTs, black points represent on-trail CTs. Mixed CTs are off-trail cameras that became on-trail when a new section of the trail was built directly adjacent to the off-trail site. On the base map, green represents forested areas, blue represents bodies of water, light blue represents glaciers, and white and beige represent bare rock and snow in high alpine areas. The inset map shows a portion of the provinces of British Columbia (BC) and Alberta (AB) with major nearby cities in white, other UNESCO Canadian Rocky Mountain Parks in light green, Mount Robson in dark green, and the Berg Lake Trail in red.

### Statistical analysis

To assess how black and grizzly bears use this heterogeneous recreation landscape, we conducted a three-stage analysis focused on behavioural shifts at different spatial and temporal scales. Together, these scales were intended to capture both reactive responses to immediate human presence and anticipatory responses to regular patterns of human activity. First, to assess weekly habitat use, we used a generalized linear mixed effects model (GLMM) to evaluate bear detections in relation to weekly recreation intensity and proximity to the trail. Second, we compared daily (diel) activity patterns and overlap between bears and humans in open and closed sections of the trail. Finally, we calculated Avoidance-Attraction Ratios (AARs) to compare direct responses to hikers in open vs. closed sections of the trail. AARs were intended to capture reactionary responses to hiker events, while weekly habitat use and diel activity were intended to capture different potential anticipatory responses to general patterns of hiker behaviour. By including all three measures, we aimed to test three competing hypotheses (i.e., fear, attraction, neutrality), while also identifying the behavioural mechanisms through which they are expressed (i.e., reactionary or anticipatory).

To measure weekly habitat use, we used the count of independent detections by CT site, species and week, as a measure of habitat use through space and time, and fit a Bayesian GLMM with a negative binomial distribution. Each study week was a 7-day period from Wednesday to Tuesday, to ensure that the weekend (when human activity is typically highest) was placed centrally within each week (Fennell et al., 2023; Green et al., 2023). We used a 30-minute independence threshold for bear detections, based on a comparison of independent detection events obtained under several candidate intervals (Figure S3). The total number of events stabilized after 10 minutes with little difference up to 60 minutes; therefore, we selected the 30-minute threshold to stay consistent with similar studies (e.g., Burton et al., 2015, 2024; Procko et al., 2022). We therefore assumed that detections of the same species within 30 minutes were the same individual or part of a non-independent group (e.g., mother and cubs or a courting pair).

For the recreation predictor variable, we calculated an effort-corrected rate of independent human detections (using a 1-minute independence threshold rather than 30 to account for the high frequency of human detections) per 100 camera-trap days as a fine-scale measure of recreation intensity at the site-level. Rather than being assigned a constant near-zero value, off-trail sites were assigned the recreation rate of the nearest on-trail site to account for the zone of influence that human disturbance exerts on wildlife, which extends beyond the immediate vicinity of the trail (Boulanger et al., 2012; Dertien et al., 2021; Parsons et al., 2020; Thompson et al., 2025). To capture spatial heterogeneity in this effect, we included proximity to trail (decay transformed with a 500-meter scale parameter) both as a main effect and in interaction with recreation. We are thus assuming that the effect of recreation is spatially diffuse and affects the larger area surrounding the site and not just the immediate view of the camera trap itself. Under this assumption, an off-trail site in an open section of the park may be more strongly affected by recreation than an on-trail site in a closed section where human activity is minimal.

To control for variation in environmental conditions, we included Normalized Difference Vegetation Index (NDVI) as an index of seasonal vegetation productivity, which is closely linked to the availability of key foods (i.e., berries, roots, herbaceous plants, and primary consumers like ungulates; Falconi et al., 2022; McLellan & Hovey, 1995; Pettorelli et al., 2005, 2011; Tuck et al., 2014), and elevation, which influences movement associated with the timing of phenology (e.g., green up), snow-pack and proximity to den sites (Bowersock et al., 2023; Pigeon et al., 2014; Servheen, 1983). This is particularly relevant for grizzly bears, which often den at high elevations and track resource phenology along elevational gradients (Berman et al., 2019; Pigeon et al., 2014). We tested NDVI and elevation for multicollinearity using Pearson’s correlation index and Variation Inflation (Table S1; Figure S4). Finally, we included a random effect for CT station to account for non-independence among repeated weekly counts of detections at individual CT stations, and a first-order autoregressive term (AR1) to account for temporal autocorrelation in residuals across consecutive weeks within camera trap sites (Shermeister et al., 2024; Zuur et al., 2009).

We built these models in the *brms* package in R, using weakly informative priors (Bürkner, 2017; R Core Team, 2024). We used a normal prior centered at 0 with a standard deviation of 2 on the fixed effects to prevent overfitting while remaining non-restrictive. On the intercept, we set a less informative normal prior centered around 0 with a standard deviation of 10 to provide mild regularization (Gelman et al., 2013). For the standard deviation of the site-level random effect, we set a half-Cauchy prior centered at 0 with a standard deviation of 2 to avoid extreme values (Gelman, 2006). Finally, we set an exponential prior (rate =1) on the negative binomial shape parameter to prevent extreme over-dispersion (Gelman, 2006). Other parameters used the default flat priors within *brms* (Bürkner, 2017). We assessed model convergence using the Rubin-Gelman statistic (Rhat>1.10; Gelman & Rubin, 1992) and inspection of trace and posterior predictive plots (Hobbs & Hooten, 2015). All models showed satisfactory convergence (Figure S6; AB7). Lastly, we performed Moran’s I tests for spatial autocorrelation. Generally, we interpreted a parameter estimate as having strong evidence of an effect if the 95% credible intervals were not overlapping 0, and weak evidence if the 80% credible intervals were not overlapping 0 (Fennell et al., 2023; Granados et al., 2023; Kéry & Royle, 2020).

To estimate daily activity patterns, we fit kernel density functions to time-of-detection data for each bear species and for humans detected at on-trail CT sites in the open and closed sections of the trail (Anderson et al., 2023; Rowcliffe et al., 2008; Wrazej, 2024). CT sites located within the mixed closure status section (Figure 1) varied in their closure status over time. Rather than being assigned a single open or closed designation for the entire study period, particularly in 2024, when areas were closed and reopened, and sections of the trail were rerouted. All images of bears and humans were included regardless of group size and without applying time-to-event filters (Peral et al., 2022; Wrazej, 2024). Each data point, therefore, represented a discrete moment when at least one bear or hiker was active at a given camera trap site. Detection times were converted to solar time using site-specific latitude and longitude to standardize observations to the local light cycle (Frey et al., 2017; Ridout & Linkie, 2009). Differences in overall activity curve shape between bears (in open and closed sections) and humans were first tested using Wald’s tests with statistical significance evaluated at α = 0.05 (Green et al., 2023; Wrazej, 2024). Temporal overlap in activity between groups (i.e., humans as well as each species of bear, in open and closed sections) was then quantified using the overlap coefficient (Δ), where Δ = 1 is complete overlap (i.e., no difference in activity between open and closed sections) and Δ = 0 is no overlap (i.e., complete temporal partitioning). We applied Δ₄ for sample sizes greater than 50 detections and Δ₁ for smaller samples (Meredith et al., 2024; Ridout & Linkie, 2009). Uncertainty in overlap estimates was assessed by bootstrapping detection times within each group (1,000 iterations), refitting kernel density functions for each resample, and recalculating the overlap coefficient to generate 95% confidence intervals (Ridout & Linkie, 2009; Wrazej, 2024). Non-overlapping intervals were interpreted as evidence of significant differences in activity patterns between groups (Green et al., 2022; Wang et al., 2015).

We used AARs to assess whether human presence causes direct temporal displacement or attraction within sites for bears using trails (Parsons et al., 2016). AARs calculate the time interval for a focal species detection before (T1) and after (T2) detection of a putatively interacting species, typically a predator, but in our case, humans. AAR (T2/T1) greater than 1 indicates that bears are avoiding the area after the passage of a human, and AAR less than 1 suggests that bears are attracted to the area after human passage (Parsons et al., 2016). Since human activity is independent of bear activity, T1 represents a baseline rate of human detections (Gump & Thornton, 2023; Niedballa et al., 2019). Therefore, when the time for a bear to return to a site exceeds this baseline (T2>T1), this signals avoidance of humans by bears (Gump & Thornton, 2023). Due to the discrepancy in detection rates between bears and humans, we used a modified AAR, which defines T1 as the time between a bear detection and the next human detection, and T2 (sometimes referred to as T2b) as the amount of time between the most recent human detection and the next bear detection (Dymit et al., 2025; Gump & Thornton, 2023; Naidoo & Burton, 2020). We set a minimum of 30 minutes and a maximum of seven months for consecutive interactions (i.e., detections must be within the same May-November of a given calendar year) to avoid overestimation of avoidance between years. We therefore assume that when a bear is detected after a human, it has sensed the presence of that human or group of humans (e.g., via auditory, olfactory, or visual cues; Niedballa et al., 2019), and any resulting systematic lag in bear return time within the same year is interpreted as a direct behavioural response rather than random co-occurrence (Figure S5).

To determine whether AAR values differed between open and closed sections of the trail, we then ran a second set of Bayesian GLMMs in *brms* (Bürkner, 2017). These models used log-transformed AARs with a Student-t distribution to account for occasional extreme AAR values (Gelman et al., 2013). We included a random effect for site and a fixed effect for the closure status of a site during a given detection. For these simple models, we used the default flat priors set by *brms* (Bürkner, 2017). Both models converged successfully (Table S2; Figure S8; SB9).

## RESULTS

We included data from our 42-camera trap array from July 12^th^, 2023, to June 24^th^, 2025, totalling 14,412 camera trap days of sampling effort within the snow-free months (May-November). Over this period, we recorded 205 independent detections of black bear and 99 independent detections of grizzly bear. Recreation intensity varied markedly along the trail, ranging from 165 to 16,410 humans per 100 camera trap days in open sections (mean= 5,009), and 1 to 3,079 in closed sections (mean= 383; Figure 2A).

**Figure 2.**
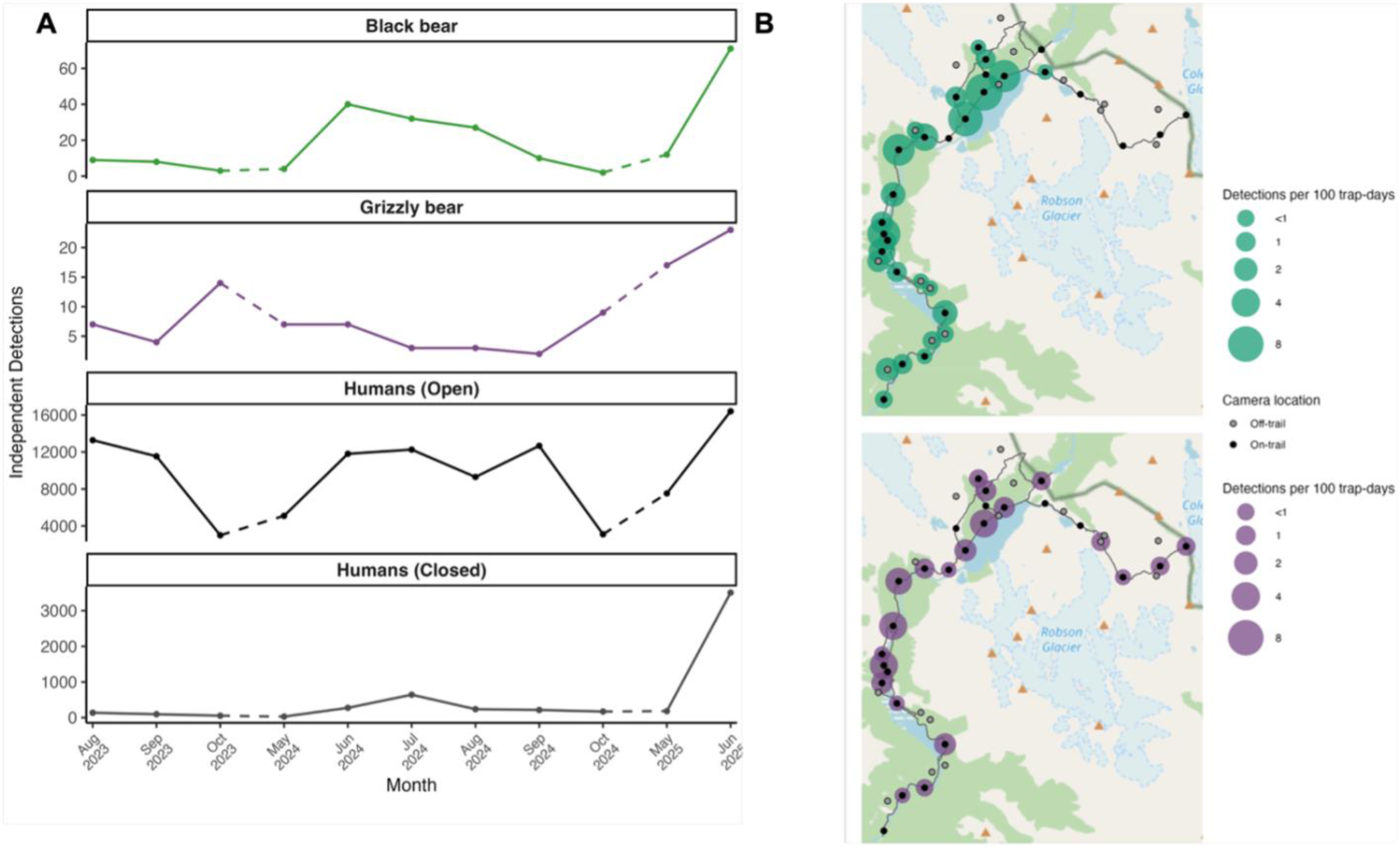
Independent monthly detection rates of black bears (*Ursus americanus*), grizzly bears (*Ursus arctos*), and humans in open and closed trail sections of Mount Robson Park, British Columbia, Canada. **(A)** Temporal patterns in detection rates from July 12, 2023, to June 24, 2025 (dashed lines indicate winter months excluded from analyses). **(B)** Spatial variation in bear detection rates across camera-trap sites over the same sampling period. Circle size represents relative detection rate (detections per 100 trap-days). Detections were considered independent using a 30-min threshold for bears and a 1-min threshold for humans.

Black bear weekly habitat use was strongly positively associated with recreation intensity and with NDVI and weakly negatively associated with elevation. Grizzly bear weekly habitat use was not associated with recreation intensity, but it was strongly positively associated with proximity to trails. Grizzly bears were also strongly negatively associated with NDVI (Figure 3). Moran’s I tests revealed no significant evidence of spatial autocorrelation for either species (Table S3). Temporal correlation structure was strong for black bears (ar[1] = 0.73 [0.27-0.99]), but not for grizzly bears (ar[1] = 0.28 [-0.87-0.97]). The random effect for CT site indicated moderate variation among sites unexplained by the fixed effects for both species (SD = 1.73 for black bears and SD = 1.22 for grizzly bears).

**Figure 3.**
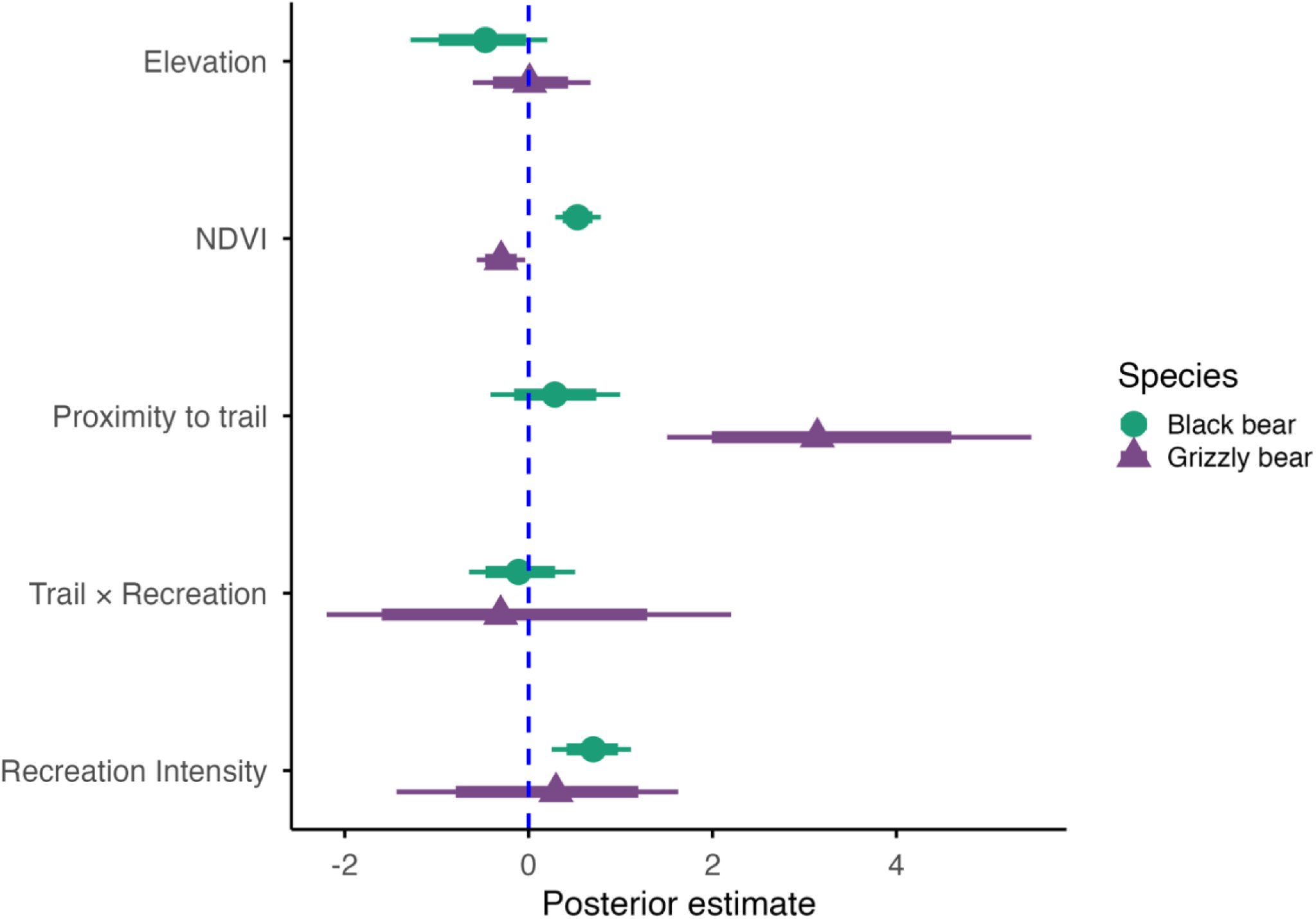
Posterior estimates (x-axis) of effects (y-axis) from the Bayesian generalized linear mixed effects models estimating weekly habitat use in Mount Robson Park, BC, Canada. Points represent posterior means, thick lines denote 80% credible intervals, and thin lines denote 95% credible intervals. Trail x Recreation represents the interaction between proximity to trail and weekly recreation. Full parameter estimates in the Appendix (Table S2).

Both black bears and grizzly bears temporally segregated their activity from humans, with both species being less active during times of heavy human trail use and exhibiting stronger avoidance of humans in open sections than in closed sections (Figure 4). In open sections, bear activity showed more pronounced peaks and troughs, whereas in closed sections, activity patterns were more even through the day. Human activity was concentrated mid-day (peaking just after solar noon), thus on open portions of the trail, bears were most active in the early morning (5:00-10:00 for black bears, and 0:00-8:00 for grizzly bears) and late afternoon (18:00-24:00 for both species), when humans were not (Figure 4a). Temporal overlap between bears and hikers was greater in closed sections than open sections for both species (i.e., non-overlapping 95% confidence intervals), but interspecific overlap was not significantly different between sections (Figure 4b). For all comparisons, the difference in temporal activity distributions was statistically significant (p<0.001).

**Figure 4.**
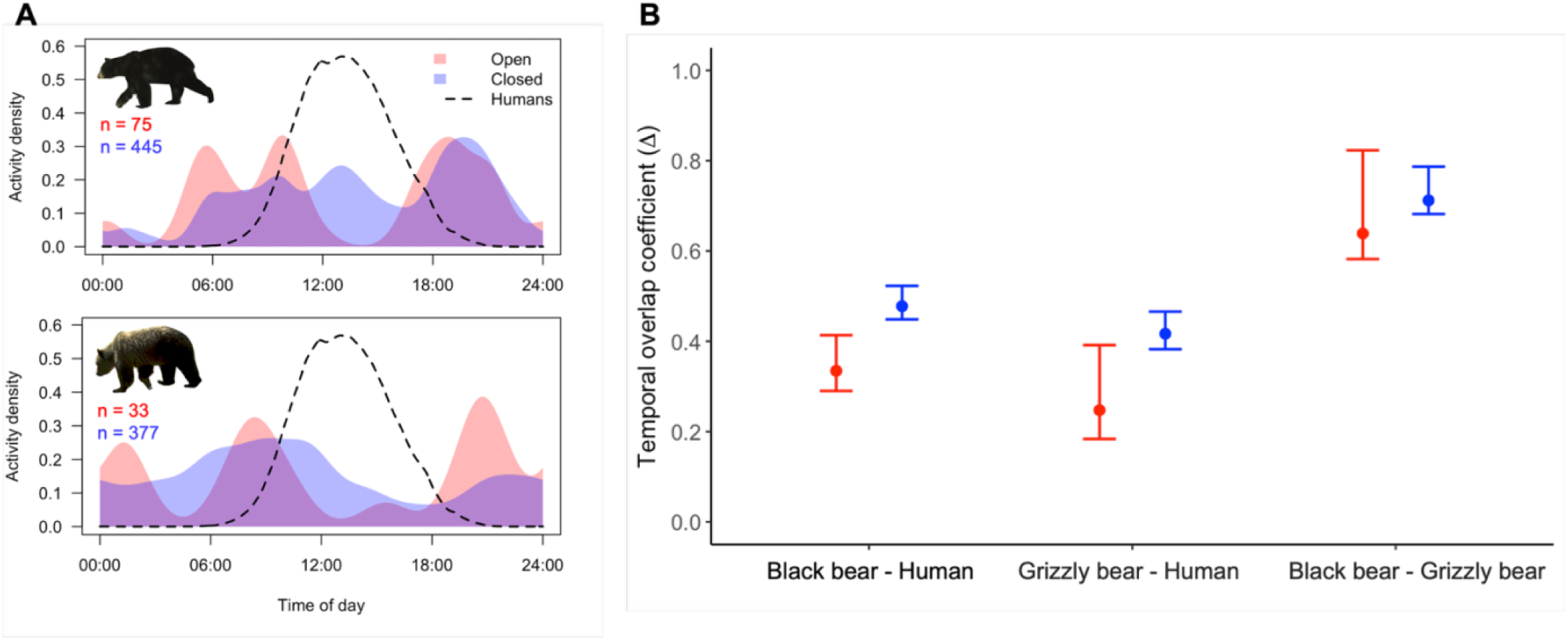
**(A)** Diel activity patterns of black bears, grizzly bears, and hikers recorded by on-trail camera traps in Mount Robson Park, BC. Shaded curves show bear activity in open (red) and closed (blue) trail sections, and the dashed black line represents hiker activity. The y-axis indicates relative activity density, and the x-axis is time of day in solar radians. Raw number of detections (n) for bears in open vs. closed sections, by species, are included in the top left corner of each. **(B)** Overlap coefficients (Δ) with 95% bootstrap confidence intervals for pairwise diel activity comparisons from on-trail camera trap detections in Mount Robson Park, BC. Red represents the comparison between bears in open sections and humans; blue represents the comparison between bears in closed sections and humans. Overlap coefficients are Δ₄ when sample size (n) ≥ 50 and Δ₁ when n < 50; uncertainty was estimated using 10,000 bootstrap iterations.

For the AAR model, we obtained 111 valid AARs (i.e., T1 and T2 < 7 months) for black bears and 56 for grizzly bears. The distribution of AAR values for both species, in both open and closed sections, was centered around 1 and thus consistent with a neutral response, indicating that the time interval between human and subsequent bear detections was similar to the interval between bear and subsequent human detections (Figure 5). The posterior distribution of the closure status effect overlapped zero, indicating no evidence of a difference in AARs between open and closed sections of the trail (Figure S8). Site-level (random effect) variance between camera trap sites was low (SD = 0.03 for black bears and 0.08 for grizzly bears), suggesting that most variation in AARs occurred within sites over time rather than between different sites.

**Figure 5.**
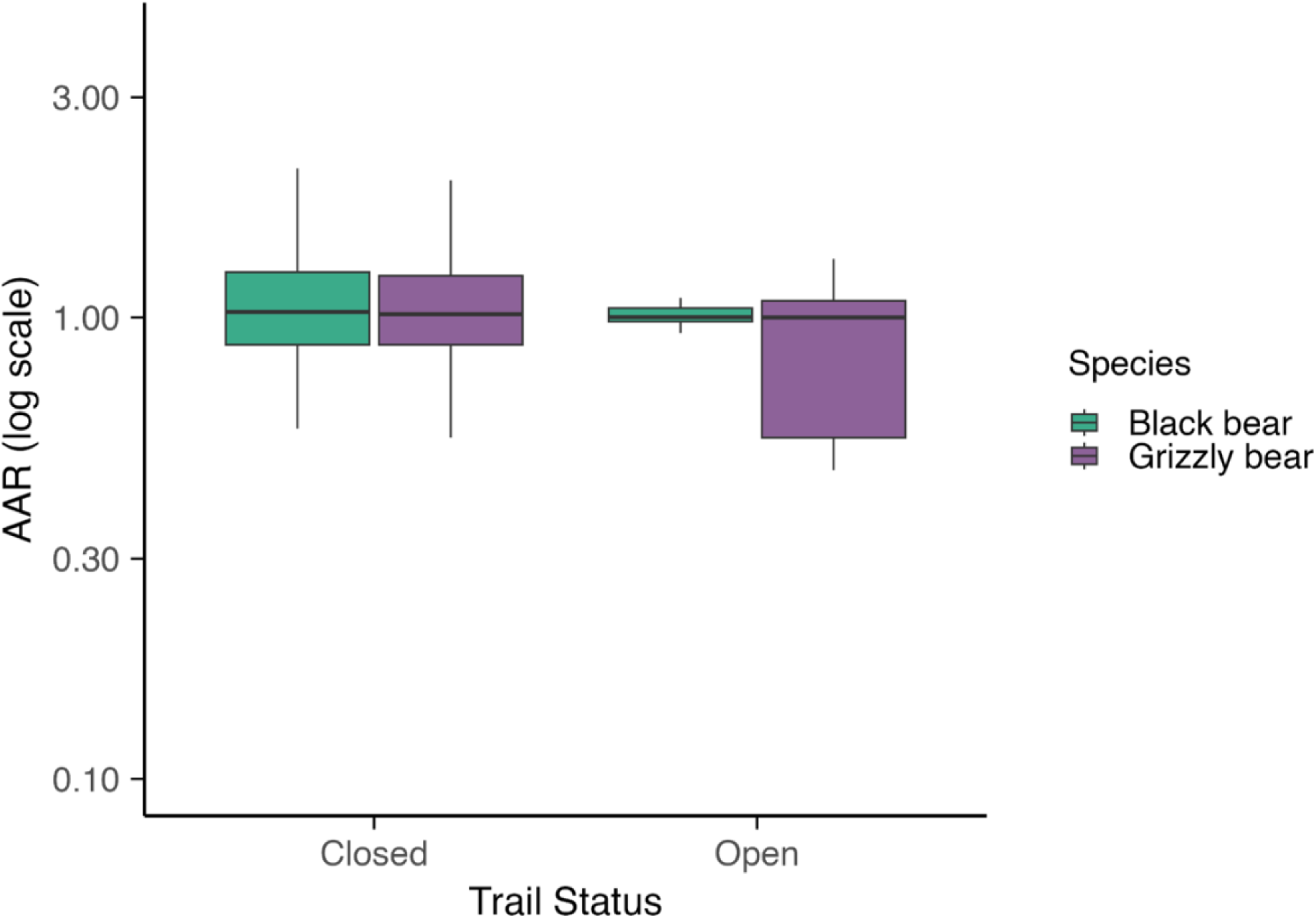
Avoidance–attraction ratios (AARs) for black bears and grizzly bears derived from camera trap detections in Mount Robson Park, BC. Boxplot shows the distribution of AAR values on the log scale, where values greater than 1 indicate avoidance of hikers, values near 1 indicate a neutral response, and values less than 1 indicate attraction. The boxes represent the interquartile range, black lines represent medians, and whiskers extend to 1.5x interquartile range.

## DISCUSSION

Across three spatiotemporal scales of bear response to recreationists (daily, weekly, and by event), black bears were detected more during high recreation weeks and sites and grizzly bears were detected more closer to trails, regardless of recreation intensity, but both species shifted to more nocturnal activity in open sections of the trail. This behaviour was consistent with both the attraction and the fear hypotheses for black bears, and the fear hypothesis for grizzly bears—although not an overwhelming fear response, given their preference for trails. These responses were also anticipatory in nature, manifesting at the weekly and daily scale, but not directly after hiker events.

Black bear responses to recreation were consistent with both the fear and the attraction hypotheses. These responses differed in scale, indicating that these hypotheses are not mutually exclusive and can even be complementary across spatiotemporal scales by allowing bears to balance resource acquisition and risk avoidance. For instance, black bears were positively associated with both NDVI and recreation at the weekly scale but avoided peak recreation hours daily. This finding suggests that the presence of humans did not limit their access to areas with more potential forage (i.e., higher NDVI) and may have even provided additional food subsidies, if the positive association was driven by attraction to garbage left behind by campers, or poorly secured food left out overnight by campers (Klees Van Bommel et al., 2022; Wheat & Wilmers, 2016). However, avoidance of peak hiking hours in open areas indicates that black bears are still associating humans with some level of risk and thus partition their use of the landscape in time to avoid direct interactions or potential conflicts (Gaynor et al., 2018; Klees Van Bommel et al., 2022).

Grizzly bears, on the other hand, showed evidence of only the fear hypothesis. Similar to black bears, they employed a daily temporal segregation strategy, avoiding peak recreation times. Their weekly detections were negatively associated with NDVI, and positively associated with proximity to trail, but were not strongly associated with weekly recreation. This pattern suggests that grizzly bears continued to use trails for ease of movement (Whittington et al., 2022), while temporally adjusting their activity to minimize the chance of encountering hikers. Past studies suggest that encounters between grizzly bears and humans tend to result in injury or death for people more often than do encounters with black bears, due to the more territorial and defensive nature of grizzlies, making access to human resource subsidies considerably more energetically costly (i.e., through stressful defense mechanisms) and risky (i.e., could lead to lethal removal; Gaynor & Green, 2026; Gunther et al., 2024; Herrero & Fleck, 1990).

While the resource-seeking hypothesis was intended to capture both food-related attraction and potential human shield effects, we found no evidence that the positive association between black bear habitat use and recreation was driven by a human shield. Grizzly bear weekly habitat use was not negatively associated with recreation, indicating there was no “shield” from the risk of intraguild predation available for black bears. Similarly, both species exhibited similar diel shifts to avoid humans; therefore, there was no evidence of black bears avoiding grizzly bears on a daily temporal scale either. We did not use AARs to compare fine-scale avoidance between the bear species, as the high number of human detections separating bear events and the potential for non-unidirectional avoidance or attraction between competing species made this approach inappropriate (Dymit et al., 2025).

In the absence of a human shield effect, our prediction was that black bear attraction to recreation activity was driven by anthropogenic resource subsidies. Synanthropic behaviours by black bears have been observed in several human-dominated systems, including urban and suburban areas where tolerance of humans may permit access to anthropogenic food subsidies (Herrero, 1983; Klees Van Bommel et al., 2022; Lewis et al., 2015; Merkle et al., 2013). To further investigate this possibility, we reviewed human-wildlife conflict data collected by MRP Park Rangers (BC Parks, 2025; Note AB1). Almost all (96%) of human-bear conflicts in the park involved black bears, which further supports the tendency of grizzly bears to avoid humans (Figure S10). Instances of people observing black bears accessing anthropogenic foods or investigating food caches, sniffing around picnic tables, and feeding on roadside trash were extremely rare (∼3% of reports). However, we found that bear activity in open areas was mostly nocturnal; therefore, the probability of visitors or staff observing these behaviours is low. While these behaviours are likely underreported, the possibility that greater black bear detections in high-recreation areas reflect food-driven attraction warrants further investigation.

In both grizzly and black bears, daily temporal segregation appeared to be the primary method for avoiding direct encounters with hikers. Since recreation in the open section is highly predictable on a daily scale (with most recreation occurring between 8 AM and 6 PM), this is a reliable strategy for minimizing risk (Gaynor et al., 2018; Kronfeld-Schor & Dayan, 2003; Lewis et al., 2021). Importantly, bears in this park had access to a large, undisturbed area that was closed to recreation, where they may seek refuge during peak recreation hours in open sections. It is therefore important to conserve refuge areas with low human disturbance in popular parks, rather than building extensive trail networks that intersect and thus fragment most of the available habitat (Lewis et al., 2021; Scallion & Titchener, 2025). Our sampling ended in June 2025, but following the park’s reopening later that month, hikers will access the full trail system, and bears will no longer have access to this low-disturbance refuge. As they are forced into more shared spaces with humans, temporal partitioning may become their only means of avoidance. Moreover, increased spatial overlap heightens the likelihood of habituation and food-conditioning, as bears may encounter human foods and garbage more frequently (Gaynor & Green, 2026). Given black bears’ preference for areas of high recreation, their greater detection rate, and the fact that most human-bear conflicts in North America involve black bears, the loss of a low-recreation refuge area may increase the risk of conflict (Penteriani et al., 2016, 2020).

There was not a significant association between trail status and AARs for either bear species, suggesting that avoidance or attraction were anticipatory rather than reactive. This finding supports the idea that bears understand patterns of human activity and are capable of shifting their use of the landscape to reduce interactions. Although recreation has been reduced in MRP over the past five years due to the trail closure, its long history of human use and proximity to the busier Jasper National Park mean bears remain accustomed to human presence and movement across the landscape. However, similar recreation studies have revealed contrasting patterns for both bear species. Notably, CT studies in Colville National Forest, Washington, USA and South Chilcotin Mountains Park, BC, Canada, where recreation rates are considerably lower, black and grizzly bears showed significant fine-scale avoidance of recreation (i.e., AAR>1; Gump & Thornton, 2023; Naidoo & Burton, 2020). These divergent results suggest that in systems where human activity is less frequent and predictable, reactionary responses may be the primary avoidance strategy for bears. In contrast, studies in other high-traffic recreation areas showed evidence of both spatial and temporal avoidance of recreationists through a shift to more nocturnal activity, or reduced use of busy areas (Burton et al., 2024; Lewis et al., 2021; Procko et al., 2023; Wheat & Wilmers, 2016). However, these trends are not consistent across all recreation areas (Fennell et al., 2023; Granados et al., 2023; Naidoo et al., 2025; Sytsma et al., 2022). This context-dependency highlights the flexibility of bear responses to human activity, which is central to understanding human-bear coexistence on a larger scale (Marion et al., 2024).

### Continuum of bear behavioural responses to recreationists

These context-dependent behavioural patterns can be interpreted within a broader continuum (Figure 6A), shaped by the frequency and predictability of human activities coupled with associated perceived risks (e.g., mortality) or rewards (e.g., anthropogenic foods). We propose that along this continuum, bear populations transition from reactive responses driven by fear (e.g., flight responses upon hearing human voices) to anticipatory responses informed by learned predictability (e.g., avoiding busy sections of a trail during the day), and eventually to neutral or habituated responses when repeated non-threatening encounters reduce perceived risk (Čapkun-Huot et al., 2024; Palmer et al., 2022; Short et al., 2024; Tablado & Jenni, 2015; Wheat & Wilmers, 2016). This continuum simplifies a range of nuanced behavioural responses and highlights broad population-level patterns among bears that continue to use the area in the presence of recreation. At the far end of the continuum, attraction is expressed when bears associate human activity with benefits (e.g., food or protection), and may arise either when habituated individuals exploit anthropogenic resources or when repeated access to such resources leads to habituation (Elfström et al., 2014; Steyaert et al., 2016; Zenth et al., 2025). Although it is rare, de-habituation can occur over time when the stimulus is removed or becomes associated with risk (Blumstein, 2016). For instance, aversive conditioning can teach habituated bears to associate humans with risk again (e.g., using loud noises or projectiles like rubber bullets), although success is highly variable and requires significant involvement from park managers that could be better allocated to preventative management strategies like visitor education (Found & St. Clair, 2018; Homstol et al., 2024; Mazur, 2010). Thus, the focus should be on preventing attraction in the first place. These responses occur along both spatial and temporal dimensions, with distinct outcomes for coexistence. In MRP, where human presence is moderately frequent and predictable for bears, anticipatory avoidance was expressed as high spatial overlap but minimal temporal overlap, the pattern likely most conducive to long-term human-bear coexistence (Figure 6B).

**Figure 6.**
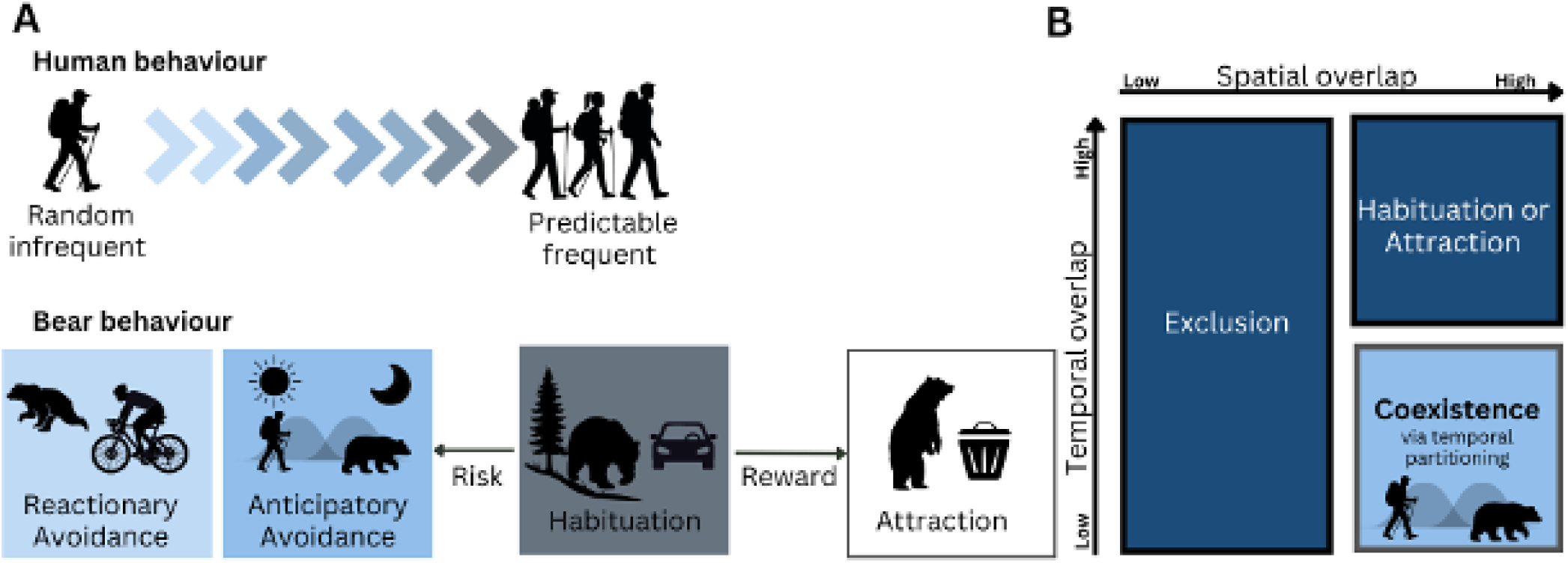
**(A)** Conceptual diagram of the continuum of bear behavioural responses to recreationists, structured around the frequency and predictability of human activity and the presence of potential rewards or risks. This figure presents a simplified representation of bear behaviour at the population level, under conditions of spatial overlap (i.e., where the population has not been displaced). Attraction is shown at the end of the continuum but may arise either when habituated bears exploit anthropogenic food resources or when repeated access to such resources leads to habituation**. (B)** Depicts the spatial and temporal dimensions of these strategies, where combinations of spatial and temporal overlap with humans yield distinct outcomes for coexistence and conservation. Dark blue denotes poor outcomes such as displacement or exclusion from protected areas or habituation leading to attraction (i.e., greater risk of food-motivated conflicts), while lighter blue denotes the outcome best aligned with human-bear coexistence. Coexistence via temporal partitioning is a form of anticipatory avoidance under which bears can access valuable areas that are shared with humans by using predictable patterns of human activity to avoid direct encounters.

Coexistence between large carnivores and humans has been defined as a dynamic state in which both humans and carnivores co-adapt to share landscapes in a manner that maintains tolerable levels of risk for both parties (Carter & Linnell, 2016). Thus, in parks and other areas where human activity is common and predictable, coexistence is best described as anticipatory avoidance by bears coupled with effective recreation management by humans. Negative interactions between bears and humans often occur when humans are present in times and places outside of what bears expect. For instance, in Yellowstone National Park, human-grizzly interactions were mostly neutral as human presence was predictable and concentrated, with most attacks occurring off-trail in backcountry areas where hikers were not expected (Gunther et al., 2024; Herrero & Fleck, 1990; MacHutchon, 2000). If coexistence is truly a process of co-adaptation, then as bears adjust their behaviour to avoid people, people must likewise adapt by maintaining predictable patterns of low-risk activities (e.g. hiking) through recreation management.

### Future directions & limitations

Some additional considerations were beyond the scope of this study but may be important when interpreting bear behavioural strategies, notably variation across seasons and individuals. During hyperphagia in the late fall, bears are so focused on food acquisition and meeting their caloric needs that they may be less risk-averse (Klees Van Bommel et al., 2020, 2022). Further, avoidance of humans may be moderated by access to resources; for instance, in a year with low berry production, we may expect bears to be more tolerant of risks associated with humans so that they can access alternative foods (Baruch-Mordo et al., 2014; Čapkun-Huot et al., 2024; Short et al., 2024; Zenth et al., 2025). Therefore, measuring and mapping both natural (e.g., berries and roots) and unnatural (e.g., human food and garbage) foods would strengthen our understanding of the costs and benefits of using human-dominated areas (Adams et al., 2017; Coogan et al., 2014, 2018; Lamb et al., 2017; Nielsen et al., 2010). Demographic variation further shapes bear responses (Kilfoil et al., 2023). Adult females with cubs tend to be more risk-averse, while young males tend to be more curious and bold (Edwards, 2023; Elfström et al., 2014; Jope, 1983; Lasky & Bombaci, 2023). We could not separately model family groups due to insufficient detections of females with cubs (Table S4), but this is a key avenue for future investigation, as differences in demographic sensitivity to recreation could have adverse effects on long-term population viability (Tablado & Jenni, 2015). Even within demographic groups, individual bears can vary markedly in behaviour and risk tolerance (Blanco et al., 2022; Bombieri et al., 2021; Lasky & Bombaci, 2023; Zenth et al., 2025). To gain a better understanding of individual variation, telemetry data (e.g., GPS collars) could complement CT sampling, providing continuous, individual-level data on movement and behaviour (Ferrer-Ferrando et al., 2023; Herraiz et al., 2024; Kröschel et al., 2017). Individual tracking could also capture changes in individual fitness or population trends (e.g., survival or reproduction), potentially revealing responses to human disturbance that may have been missed by our methods alone (Le Grand et al., 2019; Van de Walle et al., 2019). While we focused on population-level responses, we do not expect every individual within a given population to follow a uniform behavioural trajectory, as behavioural syndromes and characteristics like boldness and exploration operate at the individual level (Coppens et al., 2010; Hertel et al., 2019). We posit that anticipatory avoidance would be the prominent strategy within a given population where recreation is moderate and predictable, and the average bear has some level of experience with hikers. Studies at the individual level are critical for providing evidence on why behaviours shift, whereas population-level analyses can reveal how these changes emerge and accumulate to shape broader patterns across the population.

## MANAGEMENT IMPLICATIONS

This study offers insight into human-bear interactions in a unique setting, where bears can access a large portion of the park to spatially segregate during peak times of day. When given sufficient space, bears may choose to avoid interactions with humans. However, this pattern may shift when the Berg Lake Trail fully reopens to the public. With humans distributed along the entire trail network, bears will have reduced access to predictably safe pockets of habitat, limiting the ways they can effectively avoid humans and increasing potential conflict risk. These considerations point to the importance of hiker education and adaptive management as the park returns to high and likely growing recreation levels. Based on the greater daily temporal partitioning we observed in open areas of the trail, we recommend maintaining low human activity along trails outside of peak hours to allow bears to continue their use of the park while minimizing direct interactions with hikers through temporal segregation. This recreation management may include enforced daily closures if necessary. Further, the observed preference of grizzly bears for areas closer to trails underscores the importance of encouraging hikers to carry bear spray, especially as bears adjust to the increased human presence, in the initial years post-reopening of the trail. Given results suggesting that closed sections of the trail may have offered refuge from human activity, managers planning new or expanding trail networks should space trails in ways that preserve large, contiguous refuge areas and key habitat features (MacHutchon, 2016; Scallion & Titchener, 2025). Further, managing attractants like food and garbage is critical to ensure that these learned, proactive responses do not turn into attraction/food-conditioning. Attractant management is especially important for black bears, given their greater activity during periods of high recreation, which may increase the likelihood of encountering human foods. Effective hiker education, such as clear signage and engaging with visitors, should be implemented to encourage responsible behaviours such as remaining on designated trails, securing food and garbage, and exercising vigilance while recreating outside of peak times (Marion, 2016; Marley et al., 2017; Murphy & Halpenny, 2025; Scallion & Titchener, 2025). PAs represent some of the last remaining strongholds for large carnivores, yet even within them, human activity is pushing carnivores to the margins (Ripple et al., 2014; Venter et al., 2016). Addressing this tension is central to achieving the conservation goals of PAs. This context in MRP, where human-bear coexistence has been supported through the availability of spatial refuge and predictable patterns of human activity, may serve as a model for reconciling conservation and recreation objectives in other PAs, reminding us that large carnivore conservation and sustainable use of the ecosystem services provided by parks are not mutually exclusive.

**Table 1.** Predictor variables included in the Bayesian generalized linear mixed effects models (GLMMs) used to assess the effect of recreation on weekly habitat use for black and grizzly bears using camera trap detections in Mount Robson Park, BC, Canada.

| Variable | Description | Rationale | Mean | Min | Max |
| --- | --- | --- | --- | --- | --- |
| Recreation Intensity<sup>1</sup> | Weekly total independent human detections at each site per 100 CT days. Off-trail sites assigned recreation intensity of nearest on-trail site. | Captures spatial and temporal variation in human presence across the landscape. | 794 | 0 | 17,000 |
| Proximity to trail<sup>2</sup> | Distance in meters of a CT site to the nearest hiking trail with an exponential decay ($e^{-distance/500m}$). | Serves as a spatial proxy for human disturbance, with effects expected to diminish with increasing distance from trails and also reflects ease of movement along linear features. | 0.85 | 0.26 | 1.0 |
| NDVI<sup>3</sup> | Normalized Difference in Vegetation Index per week at each CT site, measured in 16-day intervals (250-meter resolution). Consecutive weeks may have the same NDVI value. | Serves as a proxy for primary productivity, and therefore forage availability. | 0.36 | -0.3 | 0.99 |
| Elevation<sup>4</sup> | Elevation above sea level in meters (150m resolution). | Captures seasonal elevational movements related to snowpack, temperature, phenology, and proximity to den sites. | 1576 | 842 | 2,431 |
| Effort<sup>1</sup> | Total number of active CT days within each 7-day survey week at each CT site (log transformed in the model as an offset term). | Controls for differences in detection rates caused by differences in sampling effort due to camera inactivity. | 1.0 | 6.96 | 7.0 |

1. Derived from CT sampling in this study
2. *Trail shapefiles provided by BC Parks* 3. *MODIS13Q1 (NASA) https://modis.gsfc.nasa.gov/data (Tuck et al., 2014)* 4. *extracted from the AWS Terrain Tiles DEM using the elevatr package in R (Hollister et al., 2023)*

## ACKNOWLEDGEMENTS

We would first like to acknowledge with gratitude and humility that this work takes place on the ancestral, unceded and traditional territories of the Simpcw, Lheidli T’enneh, and Lhtako Dene Nation First Nations, who have stewarded these lands since time immemorial. We thank members of the BC Parks team (including E. Ingles, A. Batho, M. Gilbert, B. May, D. Spagrud, and S. Vermeulen) for their assistance with fieldwork planning and implementation, and for contributing human-wildlife conflict data (E. Ingles and M. Gilbert). We also thank those who assisted with camera trap deployments, data collection and processing: M. Applebaum, H. Bates, A. Beatriz Pereira, L. Brennan, T. Brush, B. Carlisle, V. Duthie, E. Ferry, M. Haanen, L. Griffin, Z. Konanz, E. Tattersall, M. Wrazej, and L. Watson. This work was supported by funding from BC Parks, the Government of British Columbia, the University of British Columbia Faculty of Forestry and Environmental Stewardship, The British Columbia Chapter of the Wildlife Society, and the Natural Science and Engineering Research Council of Canada (NSERC).

## ETHICS STATEMENT

Ethics approval for this study was granted under UBC Animal Care Certificate #A25-01014 and UBC Behavioural Research Ethics Certificate #H21-01424.

