## Supporting Information for "Sharing the trail: recreation effects on bear behaviour in a Canadian Rocky Mountain Park"

April 2, 2026

Archived code and data: DOI: 10.5281/zenodo.19241617

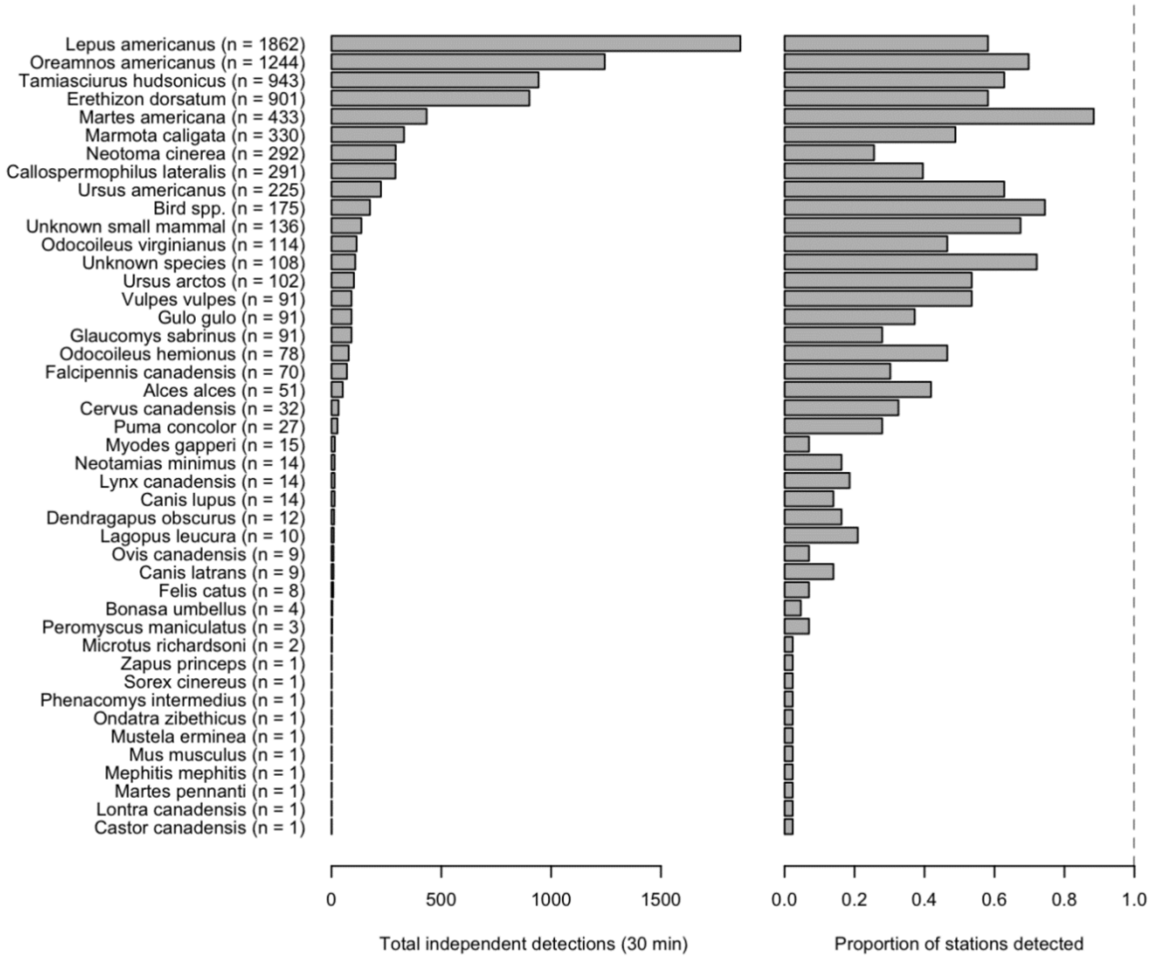

**Figure S1.** Total independent detections by species and proportion of stations detected from July 2023-June 2025, excluding humans and pets (*Canis familiaris* and *Felus catus*). A detection was considered independent if it occurred more than 30 minutes before or after another detection of the same species.

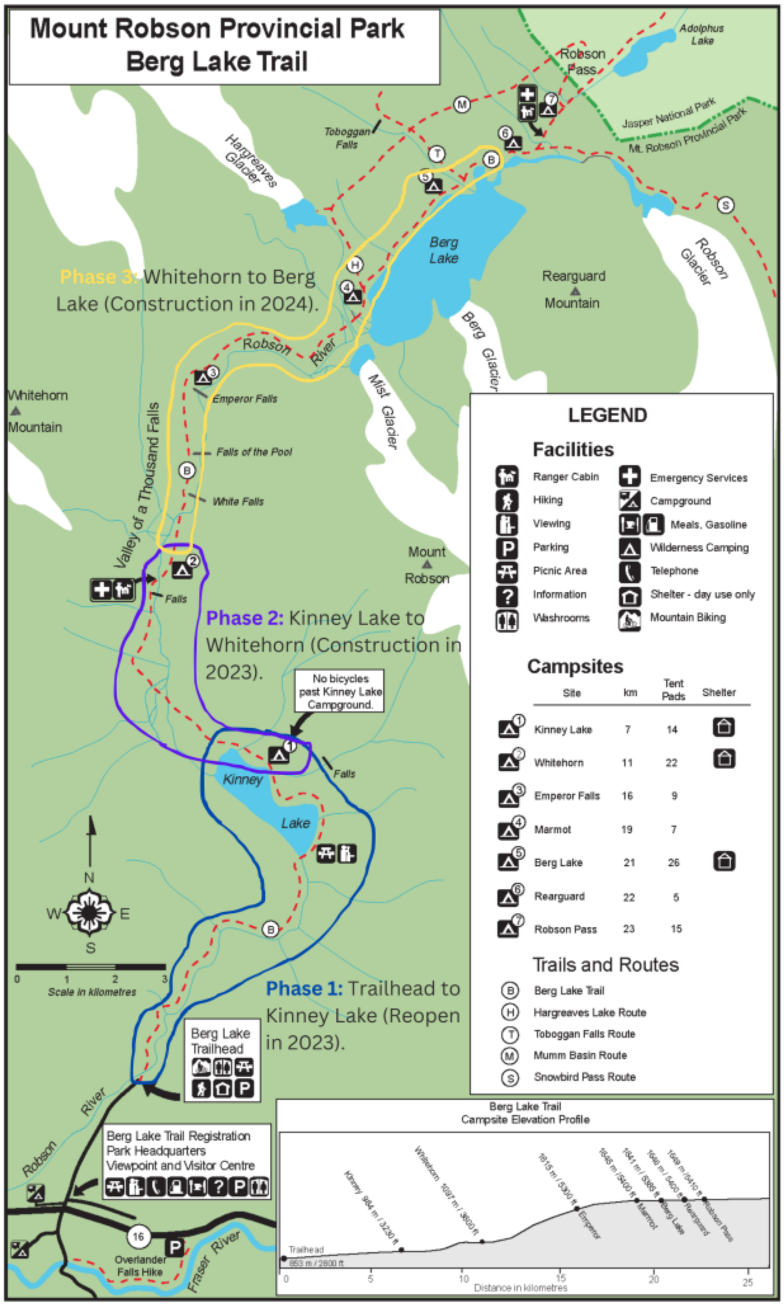


**Figure S2.** Phased reopening plan for the Berg Lake Trail (BC Parks, 2023)

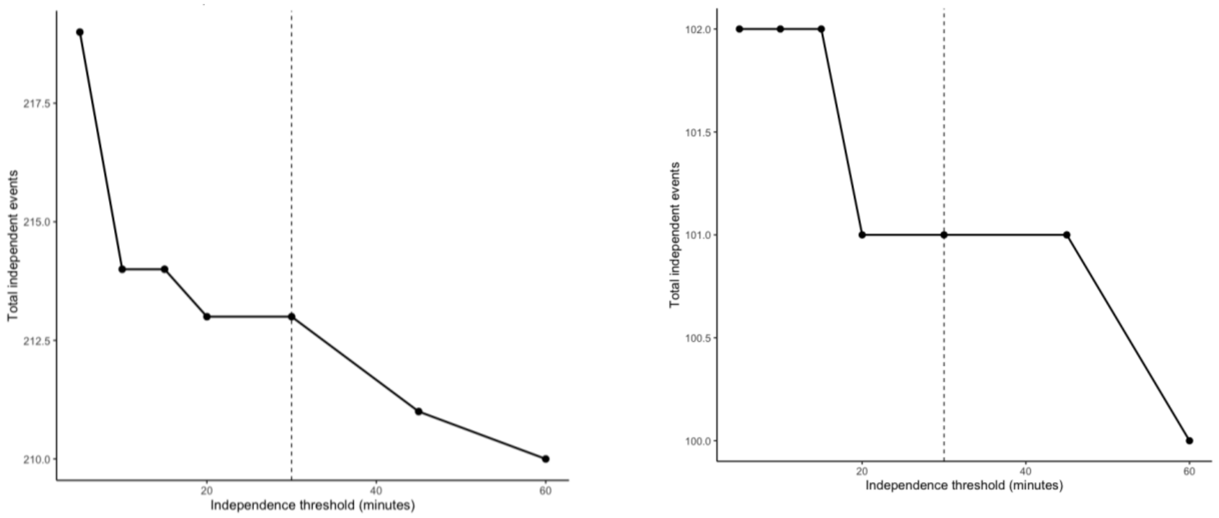


**Figure S3.** Number of total independent events under several candidate independence thresholds (5, 10, 15, 20, 30, 45 and 60 minutes) for black bear (right) and grizzly bear (left). The dashed vertical line at 30 minutes indicates the selected threshold used for all subsequent analyses.

**Table S1.** Variation Inflation Factor for predictor variables included in the weekly habitat use GLMMs, to test for multicollinearity. VIF<3 is generally considered acceptable for ecological models.

| Variable | VIF | 95% CI | Tolerance | Tolerance 95% CI |
|---|---|---|---|---|
| Proximity to trail | 1.21 | 1.17-1.26 | 0.83 | 0.79-0.85 |
| Weekly Recreation | 1.54 | 1.48-1.61 | 0.65 | 0.62-0.67 |
| NDVI | 1.42 | 1.37-1.48 | 0.70 | 0.68-0.73 |
| Elevation | 1.72 | 1.65-1.80 | 0.58 | 0.56-0.61 |

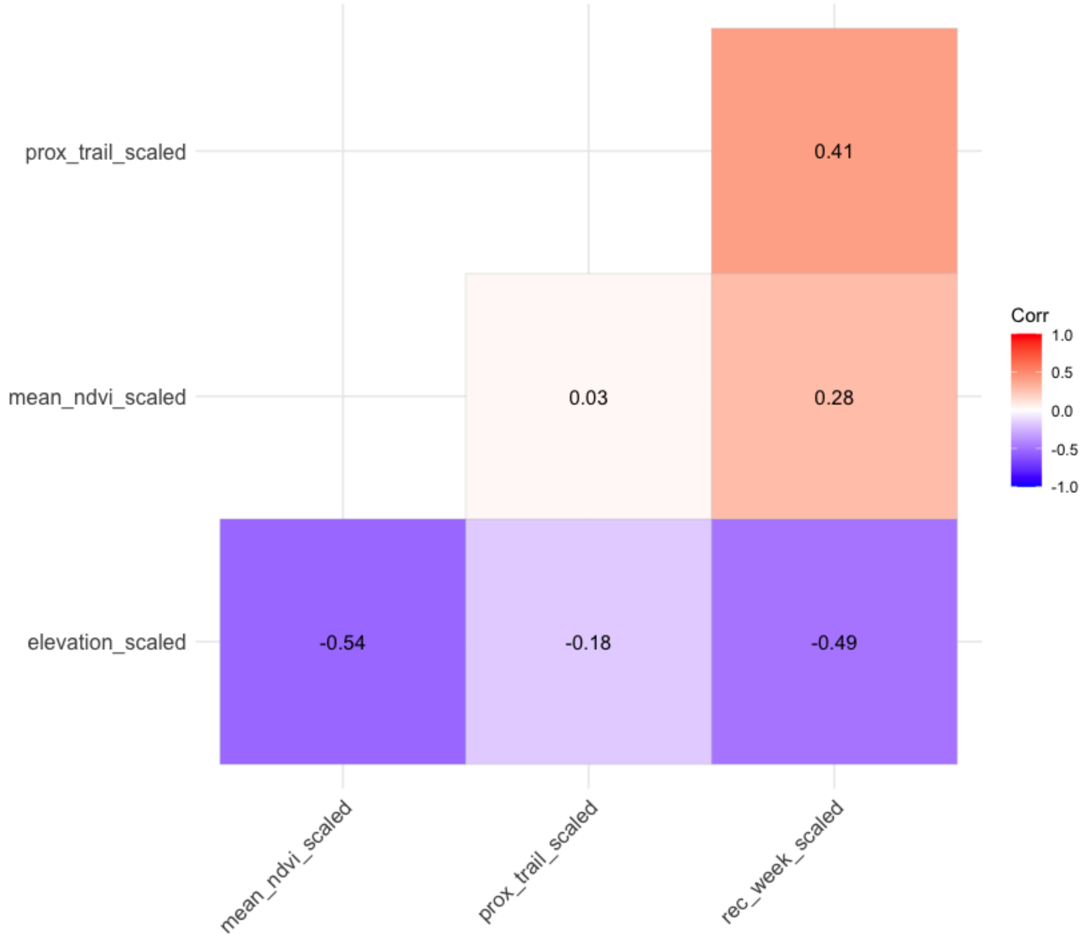


**Figure S4.** Pearson's correlation coefficient for predictor variables included in habitat use GLMM. We considered correlation |<0.6| to be acceptable.

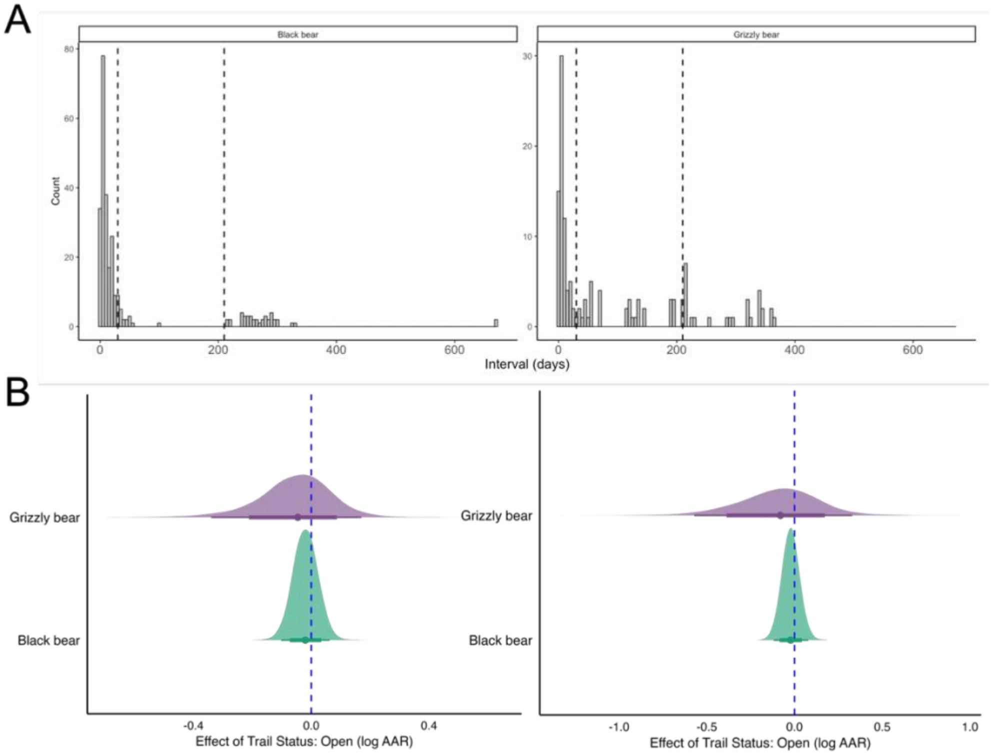


**Figure S5 (A)** Interval lengths (T1 and T2, in days) for black bears (left) and grizzly bears (right), with dashed lines indicating potential maximum thresholds of one month (30 days) and seven months (212 days). (**B**) Posterior estimates for the effect of trail status on AAR from a sensitivity analysis using each maximum interval threshold (one month [right] and seven months [left]). Points represent posterior means, thick lines denote 80% credible intervals, and thin lines denote 95% credible intervals. Posterior estimates were similar across thresholds; however, uncertainty increased with the shorter threshold due to a reduced sample size (i.e., fewer valid AARs). Because AARs are generally interpreted as fine-scale behavioural responses (Gump & Thornton, 2023; Naidoo & Burton, 2020), we conducted this sensitivity analysis to evaluate whether shortening the within-season maximum interval altered inference. Results were robust to threshold choice, supporting the use of a seven-month cutoff to maximize sample size while retaining consistent estimates.

**Table S2.** Full posterior estimates with credible intervals for the Bayesian GLMM assessing the effect of recreation and environmental variables on weekly habitat use, by species.

| Species | Term | Term_raw | Median | Lower 95% CI | Upper 95% CI | Lower 80% | Upper 80% | Rhat |
|---|---|---|---|---|---|---|---|---|
| Black bear | Elevation | b_elevation_scaled | -0.47 | -1.27 | 0.20 | -0.96 | -0.03 | 1.00 |
| | NDVI | b_mean_ndvi_scaled | 0.53 | 0.29 | 0.79 | 0.370 | 0.69 | 1.00 |
| | Proximity to trail | b_prox_trail_scaled | 0.28 | -0.40 | 0.99 | -0.16 | 0.74 | 1.00 |
| | Trail x Recreation | b_prox_trail_scaled: rec_week_scaled | -0.11 | -0.64 | 0.52 | -0.46 | 0.29 | 1.00 |
| | Weekly Recreation | b_rec_week_scaled | 0.70 | 0.25 | 1.11 | 0.41 | 0.97 | 1.00 |
| Grizzly bear | Elevation | b_elevation_scaled | 0.01 | -0.61 | 0.66 | -0.39 | 0.43 | 1.00 |
| | NDVI | b_mean_ndvi_scaled | -0.230 | -0.57 | -0.03 | -0.47 | -0.12 | 1.00 |
| | Proximity to trail | b_prox_trail_scaled | 3.13 | 1.49 | 5.40 | 1.97 | 4.57 | 1.00 |
| | Trail x Recreation | b_prox_trail_scaled:rec_week_scaled | -0.32 | -2.19 | 2.22 | -1.560 | 1.29 | 1.00 |
| | Weekly Recreation | b_rec_week_scaled | 0.31 | -1.44 | 1.61 | -0.80 | 1.19 | 1.00 |

**Table S3.** Moran's I test of spatial autocorrelation for the weekly habitat use GLMM. Spatial autocorrelation was inferred when Moran's I differed from zero and the test was statistically significant ($p < 0.05$).

| Species | Test Statistic | P-value |
|---|---|---|
| **Black bear** | -0.03 | 0.52 |
| **Grizzly bear** | -1.10 | 0.80 |

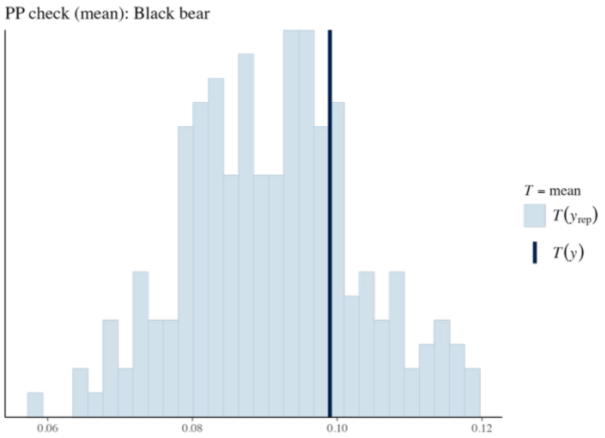


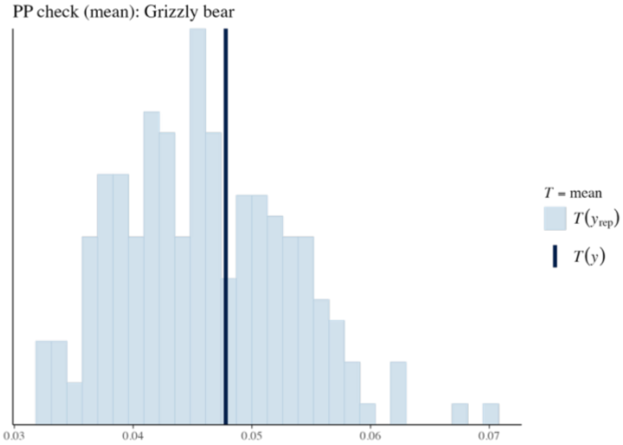

**Figure S6.** Posterior predictive check plots for weekly habitat use GLMMs, by species. Plots compare the observed mean detection rate to the distribution of simulated means from 1,000 posterior predictive draws. T(*yrep*) (light blue) is the distribution of the simulated means and T(*y*) (dark blue) is the observed mean of the actual data.

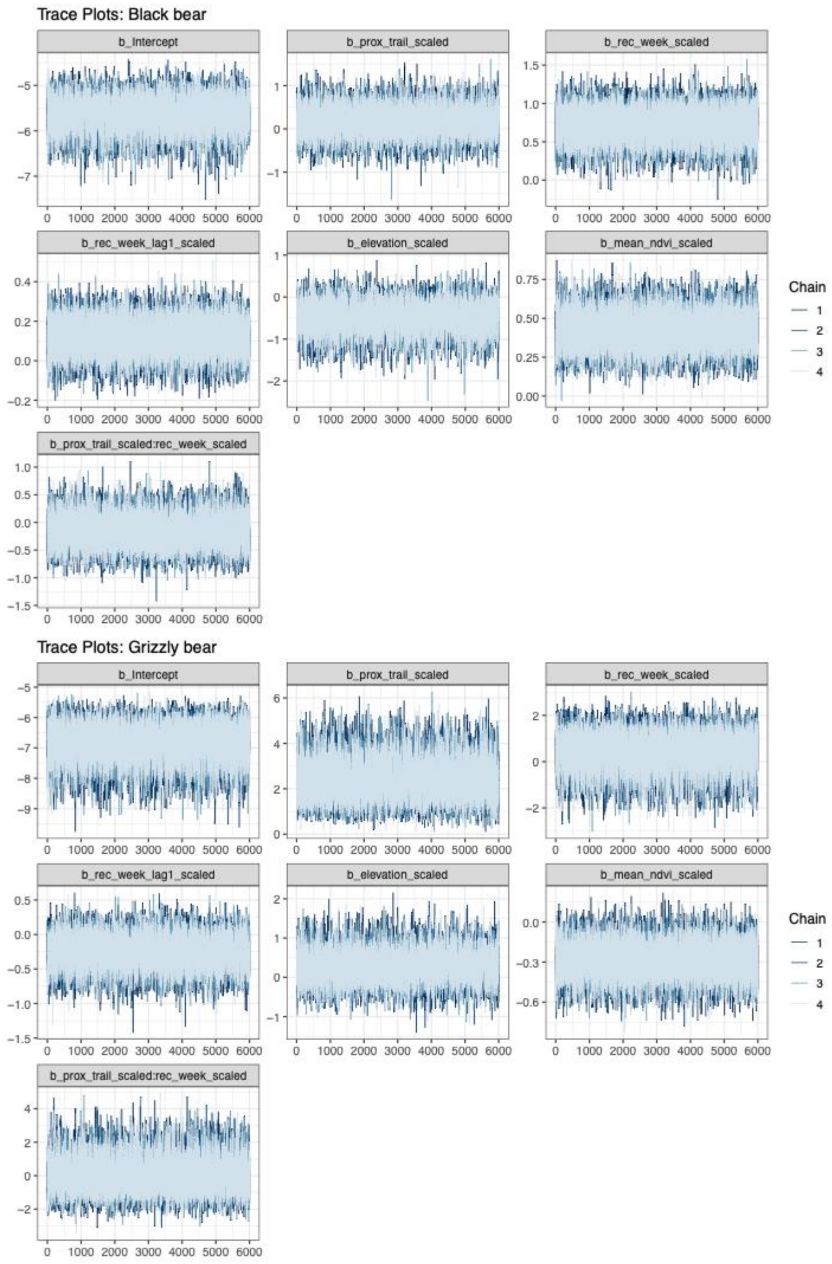

**Figure S7**. Trace plots for the habitat use GLMMs, by species. Each plot shows the Markov Chain Monte Carlo samples across four chains to assess convergence.

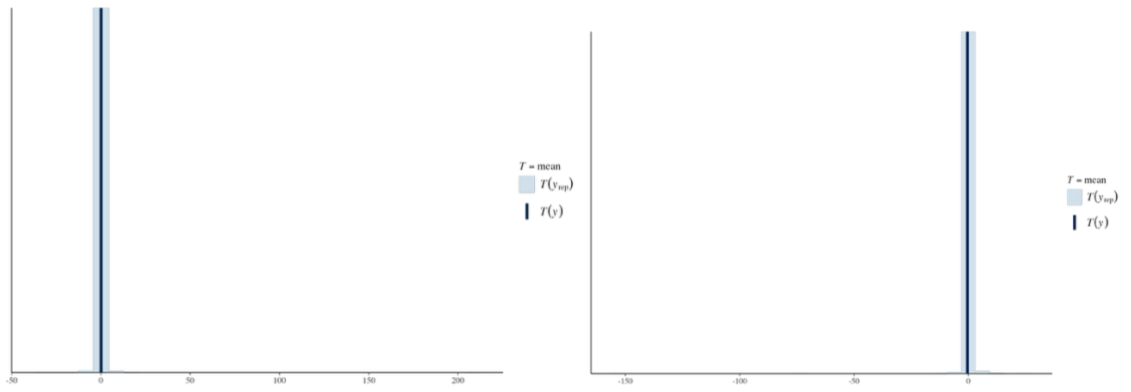


**Figure S8.** Posterior predictive check plots for AAR GLMMs, by species (black bear on the left, and grizzly bear to the left). Plots compare the observed mean detection rate to the distribution of simulated means from 1,000 posterior predictive draws. T(*yrep*) (light blue) is the distribution of the simulated means and T(*y*) (dark blue) is the observed mean of the actual data.

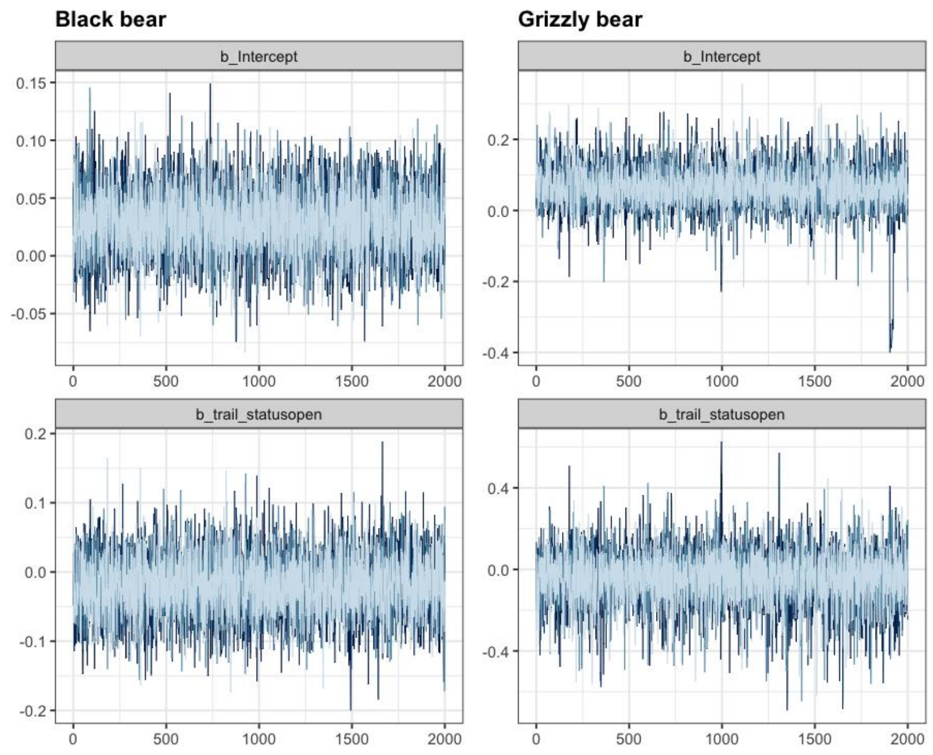


**Figure S9.** Trace plots for the AAR GLMMs, by species. Each plot shows the Markov Chain Monte Carlo samples across four chains to assess convergence.

**Table S4.** Total number of independent detections for black bears (*Ursus americanus*) and grizzly bears (*Ursus arctos*), with detections summarized by season relative to torpor and counts of observed family groups.

| Species | Total independent detections | Before torpor (September-November) | After torpor (May-August) | Family groups |
|---|---|---|---|---|
| **Black bear** | 205 | 19 | 186 | 30 |
| **Grizzly bear** | 99 | 31 | 68 | 10 |

**Note S1. Human-Wildlife Conflict Reports from BC Parks**

BC Parks Park Rangers recorded instances of human-wildlife interactions reported by staff and park users from June 1st, 2021, to September 8th, 2025, across the entire park (i.e., including other backcountry trails, frontcountry areas, and highways). For each report, managers estimated a level of habituation based on observed bear behaviours and whether the individual was repeatedly observed (Gilbert, M., *personal communication*). Surprisingly, the proportion of bears reported as tolerant of humans but not a threat (level 4) was greatest during 2022 (full closure) and 2023 (phase 1 reopening), but bears showing repeated interest in people and facilities (level 5) were only reported in 2024 and 2025, when a larger portion of the trail was open (Figure S10). We expected reports of human-bear conflicts to be considerably lower in 2022 and 2023 due to reduced overall human presence in the park; however, the reduced human activity and resulting unpredictability of those few human-bear encounters may have led to greater sightings of bears as a result of more daytime activity. The reduction in reports in 2024-2025 could be attributed to the increased nocturnality of bears under higher recreation pressure, or it could reflect reduced capacity of managers to collect and report conflict data in a fully open park. Further, some members of the public may be less likely to report sightings of large carnivores if they believe this places the animal at risk of lethal control (Martin & Burton, 2022), or if they did not perceive their interaction as a "conflict). These interpretations are largely speculative. While informative, these reports are not standardized, and likely subject to observer and reporter bias, especially the estimated level of habituation and should therefore be interpreted with caution.

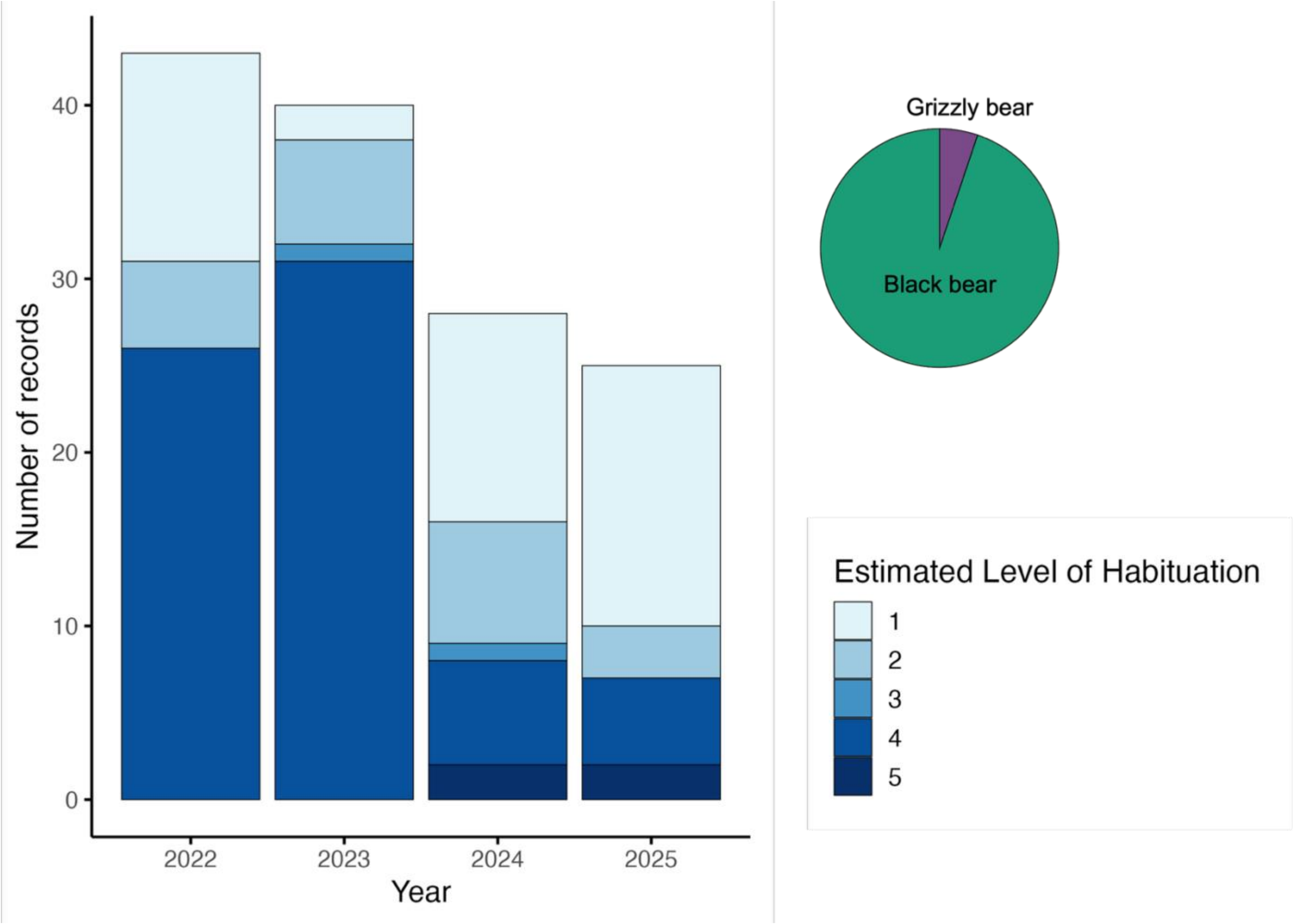


**Figure S10.** Number of human–bear conflict reports recorded by BC Parks in Mount Robson Park, BC, by year and estimated level of habituation (June 1st, 2021, to September 8th, 2025). Conflict levels were classified by park rangers as follows: 1 = Bear sighting or sign reported, 2 = Bear showing normal feeding behaviour and avoids people, 3 = Defensive aggression - Bear reacting defensively following surprise, cub(s) accompanied, cached food, and/or provoked encounter, 4 = No threat present - Bear tolerates people but ignores them and their facilities. 5 Naïve Behaviour - Bear shows repeated interest in people or their facilities, if allowed to continue, will likely result in food-conditioning or close approaches. The pie chart shows the proportion of records that involved black bears and grizzly bears.
